# The power to detect cultural transmission in the Nuclear Twin Family design with and without polygenic risk scores and in the Transmitted-Non-transmitted (alleles) design

**DOI:** 10.1101/2020.09.07.285817

**Authors:** S. Bruins, C. V. Dolan, D. I. Boomsma

**Affiliations:** Dept. of Biological Psychology, Vrije Universiteit Amsterdam, van der Boechorststraat 7, 1081 BT Amsterdam, the Netherlands, Amsterdam Public Health Research Institute, Amsterdam, The Netherlands

**Keywords:** Cultural transmission, nontransmitted alleles, genetic nurture, nuclear twin family design, power

## Abstract

The aim of our study is to compare the power of two different approaches to detect passive genotype-environment covariance originating from simultaneous cultural and genetic transmission. In the traditional Nuclear Twin Family Design cultural transmission is estimated on the basis of the phenotypic covariance matrices of the mono- and dizygotic twins and their parents, where phenotyping is required in all family members. A more recent method is the Transmitted-Nontransmitted allele design, which exploits measured genetic variants in parents and offspring to test for effects of nontransmitted alleles from parents. This design requires genome-wide data and a powerful GWA (genome-wide association) study for the phenotype in addition to phenotyping in offspring. We compared the power of both designs. Using exact data simulation, we demonstrate that compared to the Transmitted-Nontransmitted design, the Nuclear Twin Family Design is relatively well-powered to detect cultural transmission and genotype-environment covariance. The power of the Transmitted-Nontransmitted design depends on the predictive power of polygenic risk scores. Adding polygenic risk scores of realistic effect size to the Nuclear Twin Family Design did not result in an appreciable increase the power to detect cultural transmission.

---

This study’s aim is to compare the power of the Nuclear Twin Family (NTF) design and the Transmitted-Non-Transmitted alleles (T-NT) design to detect genotype – environment covariance due to cultural transmission. The classical NTF design uses the implied phenotypic covariance matrices of monozygotic (MZ) and dizygotic (DZ) twins and their parents, and the T-NT design exploits measured genetic variants in parents and offspring, to estimate genetic and cultural transmission.

Simultaneous genetic & cultural transmission leads to passive genotype-environment (GE) covariance (Plomin et al., 1977). *Passive genotype-environment covariance* occurs when parental genotypes influence the rearing environment of their offspring, which is sometimes referred to as *cultural transmission* (e.g., Cavalli-Sforza & Feldman, 1973; Eaves, 1976a; Eaves, 1976b; Eaves et al., 1977; Fulker, 1988; Maes et al., 2006). Because the offspring inherits half of each parent’s alleles, and the offspring is subject to influences of the rearing environment shaped indirectly by parents’ genotypes, a covariance arises between genotypic and environmental influences. In addition to passive GE-covariance (on which we focus in this paper), *evocative and active GE covariance* are also distinguished (Plomin et al., 1977). The latter arises when an individual’s behavior and preferences are influenced by the individual’s genotype, and the individual actively chooses and creates environments that suits their behavior and preferences. The former occurs when an individual’s actions, influences by the individual’s genotype, systematically evokes certain responses from the individual’s environment.

Genotype-environment covariance is of substantive interest as it is thought to be important in cognitive development (Cheesman et al., 2020; Scarr & McCartney, 1983; Zavala et al., 2018) and in the development of behavioral problems (Rutter & Silberg, 2002; Jaffee & Price, 2007; Bornovalova et al., 2014; Harold, et al., 2013). It is of statistical interest as the correct interpretation of model parameters in the classical twin design hinges on the assumption of no GE covariance (Keller et al., 2010). For instance, (unmodelled) covariance between A (additive genetic variable) and C (shared environmental variable) biases the estimate of the shared environmental variance (σ_C_) in the classical twin ACE model (Eaves et al, 1977; Purcell, 2002; Keller et al, 2009; Keller et al., 2010).

While there are various designs and models that allow for the estimation of genotype-environment covariance (e.g. Eaves, 1976; Carey, 1986; Dolan et al, 2014; Dolan 2020), we focus on two designs here that specifically assess cultural transmission. The NTF design (Keller et al., 2009) extends the classical twin design by including the parents of the twins. In the NTF design, the family environment is defined as an environment shared between all family members that arises due to cultural transmission. Additionally, one can estimate the variance due to sibling shared environment (environment shared between members of a twin pair / sibling pair), variance due to non-additive genetic effects, unshared environment (all variance due to influences unique to the individual, including measurement error), genotype-family environment covariance, and phenotypic assortative mating.

A more recent design to detect GE covariance stemming from cultural transmission is the transmitted-nontransmitted (T-NT) alleles design. In this design, cultural transmission is manifest in the effect on the offspring phenotype of the non-transmitted alleles (Kong et al., 2018; Bates et al., 2018). This method can be applied to parent-offspring trios, but can easily be extended to parents and multiple offspring, including twins. As parents pass on half of their alleles to their offspring, the polygenic risk score (PRS) of the offspring is a function of the *transmitted alleles* from both parents. For the other non-transmitted half of the parental alleles, a *nontransmitted PRS* can be calculated. In linear regression, the regression coefficient in the regression of offspring phenotype on the nontransmitted PRS should be zero, if genetic transmission is the only pathway of transmission from parent to offspring. Rejection of the (null) hypothesis that the regression coefficient is zero suggests that the parental genotype has an indirect effect on the offspring phenotype, i.e. that cultural transmission is present.

Our aim is to compare the power of the NTF design and the T-NT design to detect cultural transmission and to assess whether the addition of PRS to the NTF design improves the power to detect cultural transmission in this design. The outline of this paper is as follows. First, we present the classical NTFD, the transmitted-nontransmitted design, and the NTFD including PRS. Second, we outline our strategy with respect to simulation and power analysis. Third, we present the results of our simulation studies and discuss the implications.

## The Nuclear Twin Family Design

The relationship between the total additive genetic variance as inferred in the classical NTF design and the variance explained by the PRS is as follows. Suppose that there are *M* genetic variants (GVs) contributing to the variance of phenotype *ph*, and suppose that we have measured all *M* relevant GVs. Assuming for convenience (but without loss of generality) that the GVs are in gametic phase (linkage) equilibrium, we have, for individual *i*:

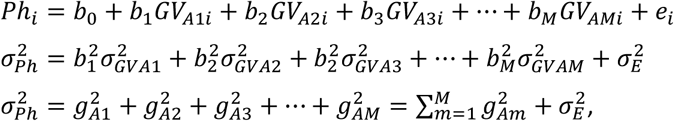

where 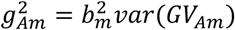, i.e., the additive genetic variance due to *GV*_*m*_, and 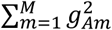 is the total additive genetic variance. The additive genetic variance as estimated in the NTF Design or the classical twin design is an estimate of 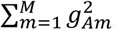, which may be biased given violations of the model (design) assumptions. Since the PRS is based on a subset of genetic variants, the additive genetic variance is the sum of the variance of the PRS based on *T* measured alleles, and the remaining latent additive genetic variance: 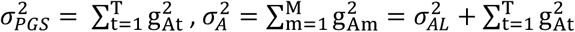, and the proportion of explained additive genetic variance is 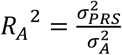. Since the total additive genetic variance 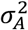 is identified by the implied covariance between MZ and DZ twins and their parents, 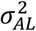 and 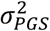 are 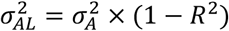 and 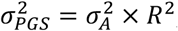, respectively.

The path diagrammatic representation of the classical NTF design, given random mating, is given in Figure 1. Using path tracing (Keller, et al., 2009), we can deduct the model-implied variances and covariances. In the NTF design, we then have

**Figure 1.**
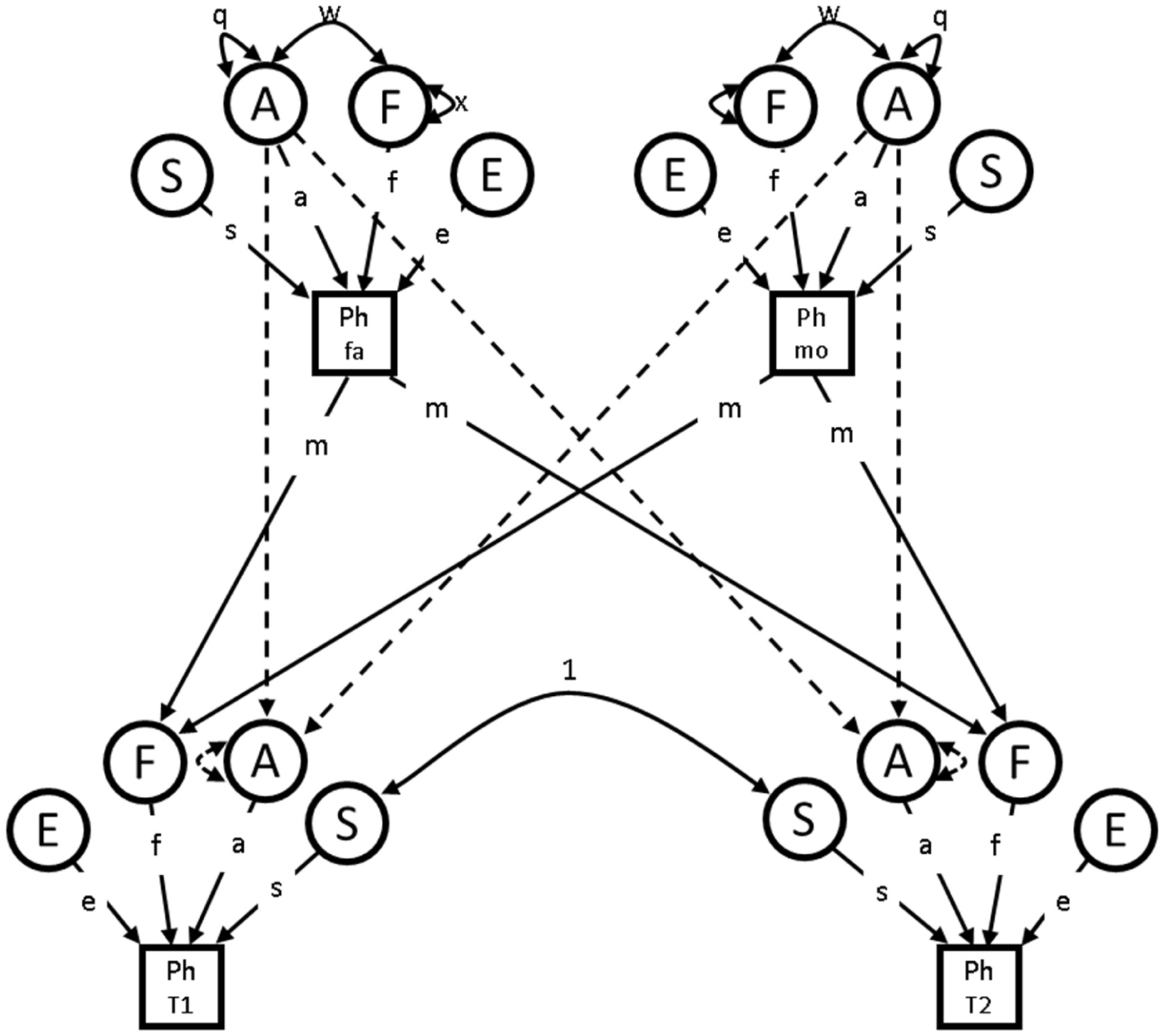
Pathdiagram of the classical Nuclear Twin Family Design given random mating (in the parameterization and notation of Keller et al., 2009). The circles denote latent variables, the squares are observed / measured phenotypic values. Single-headed arrows are paths, double-headed arrows indicate covariances. Solid lined are free parameters, dashed lines are fixed parameters. Dashed paths between parents-offspring A are fixed to .5. A = additive genetic, F = family environment due to cultural transmission, S = sibling environment shared between twins, E = unshared environment, Ph = phenotype, m = cultural transmission, w = covariance between family environment and additive genetic variable. Constraints include 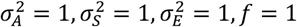.

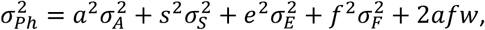

where the *q* and *x* in Figure 1 equal 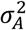 and 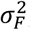, respectively. While *w* is the covariance between the latent variables *A* and *F*, the term 2*afw* is the total contribution of the covariance between genotype *A* and family environment *F* to the phenotypic variance. Given the scaling in Figure 1 (based on Keller et al, 2009), we have

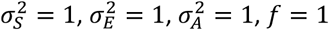, so that the equation can be rewritten as

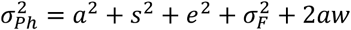

We find the variance of the family-environment 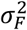 and the genotype-family environment covariance *w* by 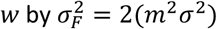 and 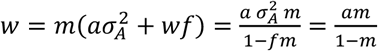, given 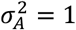 and *f* = 1. Note that *w* ≠ 0 and 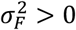 only if there is cultural transmission, i.e., *m* ≠ 0.

The Nuclear Twin Family Design including polygenic risk scores is depicted in Figure 2. Using path tracing and given the same scaling and constraints as in the classical NTFD, the complete model-implied covariance structure is

**Figure 2.**
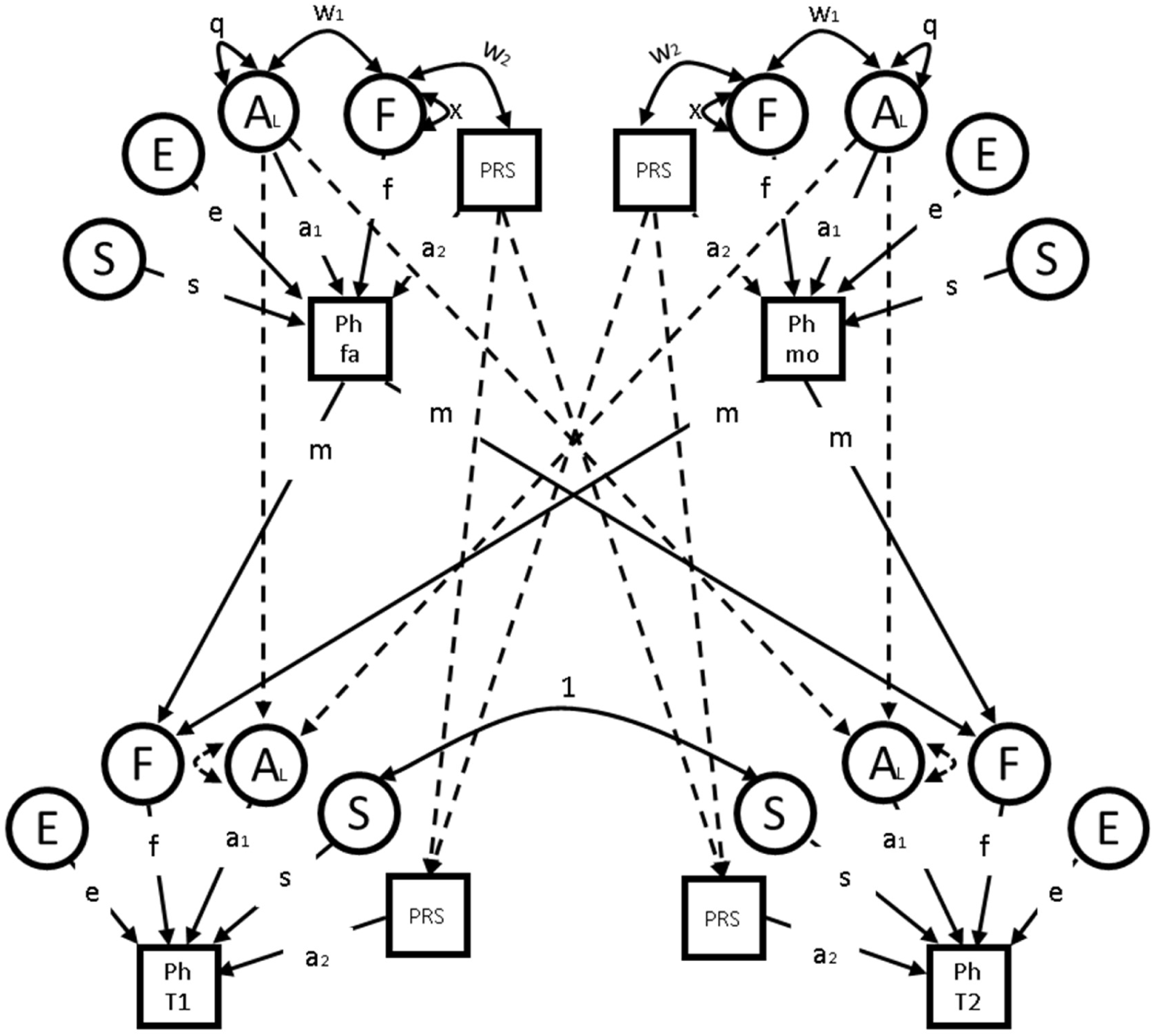
Pathdiagram of the Nuclear Twin Family Design with PRS subject to random mating. The circles denote latent variables, the squares are observed / measured values. Single-headed arrows are paths, double- headed arrows indicate covariances. Solid lined are free parameters, dashed lines are fixed parameters. Dashed paths between parents-offspring A_L_ and PRS are fixed to .5. A_L_ = latent additive genetic, PRS = observed (transmitted) additive genetic, F = family environment due to cultural transmission, S = sibling environment shared between twins, E = unshared environment, Ph = phenotype, m = cultural transmission path, w1 and w2 = covariance between family environment and latent and observed additive genetic variables. Constraints include 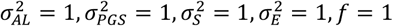.

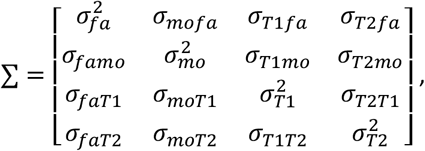

Where *fa, mo, T*1 and *T*2 stand for the phenotypic (co)variance of father, mother, twin 1, and twin 2, respectively. Variances are assumed to be equal, and (genotypic as well as cultural) transmission is assumed to be equal for both parents, such that

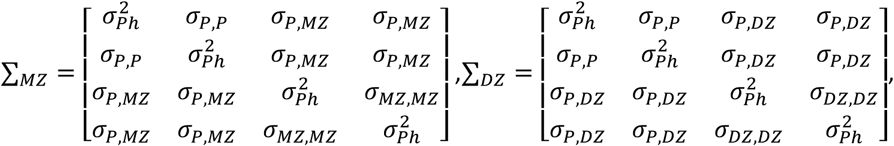

where the total phenotypic variance is 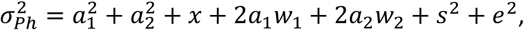 the covariance between parents is σ_*P,P*_ = *x*, the covariance between parent and (twin) offspring 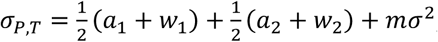, the covariance between MZ twins is 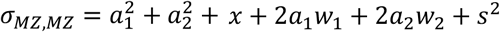, and the covariance between DZ twins is 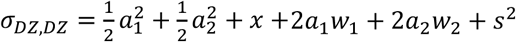 (equations and matrices adapted from Keller et al, 2009).

## The transmitted-nontransmitted PRS design

In the Transmitted-nontransmitted (T-NT) design, the phenotype is regressed on the transmitted and nontransmitted PRS, such that for individual *i*, we have

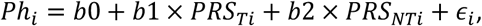

where *PRS*_*Ti*_ is the PRS that was transmitted from parents to individual *i*, and *PRS*_*NTi*_ is the nontransmitted PRS of individual *i*, based on the alleles that were not transmitted to individual *i*. While the transmitted PRS in the transmitted-nontransmitted design is equal to the PRS in the NTFD, the relation between the nontransmitted PRS in the T-NT design and the cultural transmission effects in the NTFD is more complicated. In the NTFD, cultural transmission processes are captured by the family environment, the genotype-family environment covariance, and the cultural transmission from the parental phenotype to the family environment itself. In the T-NT design, however, cultural transmission is solely represented by the regression of the offspring phenotype on the nontransmitted parental PRS.

## Power

Model identification of the NTF is well established, and the addition of PRS does not pose an identification problem. However, within an identified model, the power in tests concerning the parameters is an open question. We conducted power analysis in the NTF models using exact data simulation (van der Sluis et al., 2008). Specifically, the power detect cultural transmission was calculated as the power to reject a misspecified model in which *m* = 0, when in truth, *m* took values of *m* = .05, *m* = .10, *m* = .15, and *m* = .20. Data were simulated for various parameter settings of *a*^2^, *s*^2^, and *e*^2^ (and *m* = .05 to. 20). Detailed model parameters and variance decomposition per scenario are given in Table 1. The polygenic risk scores were simulated such that PRS explained 10% of the additive genetic variance (i.e. *R*_A_^2^ = .10). Since the noncentrality parameter is linearly related to sample size, the NCP was weighted for the number of families (*N*), and power was calculated for *N* = 100 − 10.000 (with an MZ/DZ family ratio of 1).

**Table 1.** Parameter settings and variance components for 12 data simulations.

| Sim | Par. Set. |  | Variance components |  |  |  |  |  |  |  |
| --- | --- | --- | --- | --- | --- | --- | --- | --- | --- | --- |
| | $a^2$ | $m$ | $\sigma_{AL}^2$ | $\sigma_{PRS}^2$ | $\sigma_F^2$ | $\sigma_S^2$ | $\sigma_E^2$ | $2\sigma_{AL,F}$ | $2\sigma_{PRS,F}$ | $\sigma_{Ph}^2$ |
| 1 | 0.8 | .05 | .72 (.45) | .08 (.05) | .01 (.01) | .20 (.13) | .50 (.31) | .08 (.05) | .01 (.01) | 1.59 |
| 2 | 1.0 | .05 | .90 (.50) | 0.1 (.06) | .01 (.01) | .20 (.11) | .50 (.28) | .10 (.05) | .01 (.01) | 1.81 |
| 3 | 3.0 | .05 | 2.70 (.67) | .30 (.07) | .02 (.01) | .20 (.05) | .50 (.12) | .28 (.07) | .03 (.01) | 4.04 |
| 4 | 0.8 | .10 | .72 (.42) | .08 (.05) | .04 (.02) | .20 (.12) | .50 (.29) | .16 (.09) | .02 (.01) | 1.71 |
| 5 | 1.0 | .10 | .90 (.46) | .10 (.05) | .04 (.02) | .20 (.10) | .50 (.26) | .20 (.10) | .02 (.01) | 1.96 |
| 6 | 3.0 | .10 | 2.70 (.61) | .30 (.07) | .09 (.02) | .20 (.05) | .50 (.11) | .60 (.14) | .07 (.02) | 4.46 |
| 7 | 0.8 | .15 | .72 (.39) | .08 (.04) | .09 (.05) | .20 (.11) | .50 (.27) | .25 (.14) | .03 (.02) | 1.87 |
| 8 | 1.0 | .15 | .80 (.42) | .10 (.05) | .10 (.05) | .20 (.09) | .50 (.23) | .32 (.15) | .04 (.02) | 2.15 |
| 9 | 3.0 | .15 | 2.70 (.54) | .30 (.06) | .22 (.05) | .20 (.04) | .50 (.10) | .95 (.19) | .11 (.02) | 4.98 |
| 10 | 0.8 | .20 | .72 (.35) | .08 (.04) | .17 (.08) | .20 (.10) | .50 (.24) | .36 (.17) | .04 (.02) | 2.07 |
| 11 | 1.0 | .20 | .90 (.38) | .10 (.04) | .19 (.08) | .20 (.08) | .50 (.21) | .45 (.19) | .05 (.02) | 2.39 |
| 12 | 3.0 | .20 | 2.70 (.48) | .30 (.05) | .45 (.08) | .20 (.04) | .50 (.09) | 1.35 (.24) | .15 (.03) | 5.65 |

*Note.* $\sigma_{Ph}^2$ is the total phenotypic variance given specified parameter settings. $\sigma_A^2$ , $\sigma_F^2$ , $\sigma_S^2$ , $\sigma_E^2$ , and $\sigma_{A,F}$ indicate the phenotypic variance that is explained by genotype, family environment, sibling environment, unshared environment, and genotype-environment covariance, respectively. This decomposition is based on $\sigma_{Ph}^2 = a^2 + s^2 + e^2 + \sigma_F^2 + 2aw$ . Standardized variance components are in parentheses (proportions of phenotypic variance). Par. Set. = Parameter settings. Parameter settings represent unstandardized input parameters, where $e^2 = .50$ and $s^2 = .20$ over all simulations

Given these parameter settings, we compared the power to detect cultural transmission in the NTF design and T-NT design, given identical parameter settings and sample sizes, using ordinary (i.e., not exact) data simulation. To compare power of the T-NT design with that of the NTF design, we must ensure that the sample sizes are comparable. To do so, we determined the number of families for which, in the NTFD+PRS model, the power to reject *m* = 0 was .80, given specific parameter settings. We performed a general estimation equations (GEE; e.g., Minicǎ et al. 2013) analysis on the phenotype and (transmitted and nontransmitted) PRS data of both members of twin pairs. GEE automatically adjusts the standard errors and test statistic for the dependency. Since the twins are related, the effective *N* is defined as 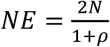, where ρ is the twin correlation. For instance, if we find that the NTFD+PRS requires 1000 twin-families (i.e. 2000 individual twins) to reject *m* = 0 with a power of .80, we use *N* = 2000 for the GEE analysis. If then, for example, *ρ*_*MZ*_ = .6 and *ρ*_*DZ*_ = .4, we effectively have 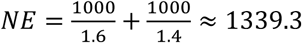 unrelated individuals. Therefore, the NCP from the GEE was corrected for the effective number of unrelated individuals.

For the NTF model, we used the NCP to calculate the power to detect cultural transmission. Given exact data simulation, the NCP equals the value of the loglikelihood ratio test statistic in the test of *m* = 0. For the simulations involving the T-NT model, we used ordinary simulation (not exact), so that the NCP is the average test statistic of the regression coefficient of the nontransmitted PRS, minus the degrees of freedom, which was obtained by running 5000 replications. The test statistic in the GEE is a robust Z-statistic. Since asymptotically, *Z*^2^ = *χ*^2^, the squared test statistics follows a (central) *χ*^2^ distribution (1 df), under the null hypothesis. Power and required sample sizes are reported given *α* = .05. Analyses were conducted in R (version 3.5.1; R Development Core Team, 2018). Structural equation modeling was performed in OpenMx (Neale et al., 2016), using the NPSOL optimizer. For the GEE modeling, the R-package *gee* was used (Carey et al., 2012). R-scripts for the simulations and model fitting are provided in the supplementary materials.

## Results

The transmitted and nontransmitted PRS are uncorrelated, and the regression coefficients of the nontransmitted PRS increased with increasing values of *m*, indicating that the simulated PRS indeed captured the cultural transmission effects. As expected, the regression coefficient of the transmitted PRS also increases with increasing *m*, reflecting the presence of cultural transmission. Model information and power calculations for the three models are given in Table 2. As can be seen from Table 2, the power to reject *m* = 0 does not differ substantially between the classical NTFD and the NTFD with PRS, but is the NTFDs have greater power to reject *m* = 0 than the T-NT design. *Figure 3* displays the power to detect the cultural transmission effects of scenario 7 over a range of sample sizes (N= 100-10000).

**Table 2.** Required sample sizes and power for NTF design, NTF design+PRS, and T-NT design.

|  |  | NTFD |  |  |  |  |  | T-NT |  |  |  |  |  |  |  |  |
| --- | --- | --- | --- | --- | --- | --- | --- | --- | --- | --- | --- | --- | --- | --- | --- | --- |
|  |  | NTFD |  |  | NTFD + PRS |  |  | Transmitted |  |  |  | Nontransmitted |  |  |  |  |
| Sim | nfam | -2LL | p | pow | -2LL | p | pow | NE | b (s.e.) | Z | p | b (s.e.) | Z | p | pow | .80 |
| 1 | 3500 | 7.4 | .006 | .78 | 7.9 | .005 | .80 | 4515 | .30 (.02) | 17.65 | <.001 | .01 (.02) | 0.85 | .397 | .11 | 50369 |
| 2 | 3100 | 7.3 | .007 | .77 | 7.8 | .005 | .80 | 3937 | .33 (.02) | 17.34 | <.001 | .02 (.02) | 0.88 | .395 | .14 | 40100 |
| 3 | 2000 | 7.1 | .008 | .76 | 7.8 | .005 | .80 | 2395 | .58 (.04) | 16.03 | <.001 | .03 (.04) | 0.81 | .403 | .13 | 27924 |
| 4 | 800 | 7.2 | .007 | .77 | 7.8 | .005 | .80 | 1011 | .31 (.04) | 8.59 | <.001 | .03 (.04) | 0.85 | .400 | .12 | 10359 |
| 5 | 720 | 7.2 | .007 | .77 | 7.8 | .005 | .80 | 896 | .35 (.04) | 8.52 | <.001 | .04 (.04) | 0.85 | .396 | .14 | 9403 |
| 6 | 480 | 7.3 | .007 | .77 | 7.9 | .005 | .80 | 564 | .61 (.08) | 7.95 | <.001 | .06 (.08) | 0.79 | .413 | .12 | 7100 |
| 7 | 330 | 7.2 | .007 | .77 | 7.8 | .005 | .80 | 408 | .33 (.06) | 5.65 | <.001 | .05 (.06) | 0.86 | .398 | .12 | 4209 |
| 8 | 300 | 7.2 | .007 | .77 | 7.8 | .005 | .80 | 365 | .37 (.07) | 5.63 | <.001 | .06 (.07) | 0.86 | .397 | .14 | 3715 |
| 9 | 180 | 7.3 | .007 | .77 | 7.8 | .005 | .80 | 207 | .64 (.13) | 4.97 | .001 | .10 (.13) | 0.76 | .419 | .13 | 2460 |
| 10 | 170 | 7.4 | .006 | .78 | 7.9 | .005 | .80 | 205 | .35 (.09) | 4.13 | .005 | .07 (.09) | 0.83 | .399 | .13 | 2044 |
| 11 | 150 | 7.4 | .006 | .78 | 7.9 | .005 | .80 | 179 | .39 (.10) | 4.03 | .006 | .08 (.10) | 0.81 | .401 | .14 | 1868 |
| 12 | 90 | 7.5 | .006 | .78 | 7.9 | .005 | .80 | 102 | .68 (.19) | 3.58 | .015 | .13 (.19) | 0.70 | .424 | .12 | 1278 |

*Note.* Sim stands for simulation scenario, pow stands for power, -2LL is the -2 loglikelihood, Z is the Z-statistic of *b*. nfam is the number of families, NE is the effective number of unrelated individuals (rounded to the nearest integer), .80 is the effective number of unrelated individuals required for a power of .80 (rounded to the nearest integer)

**Figure 3.**
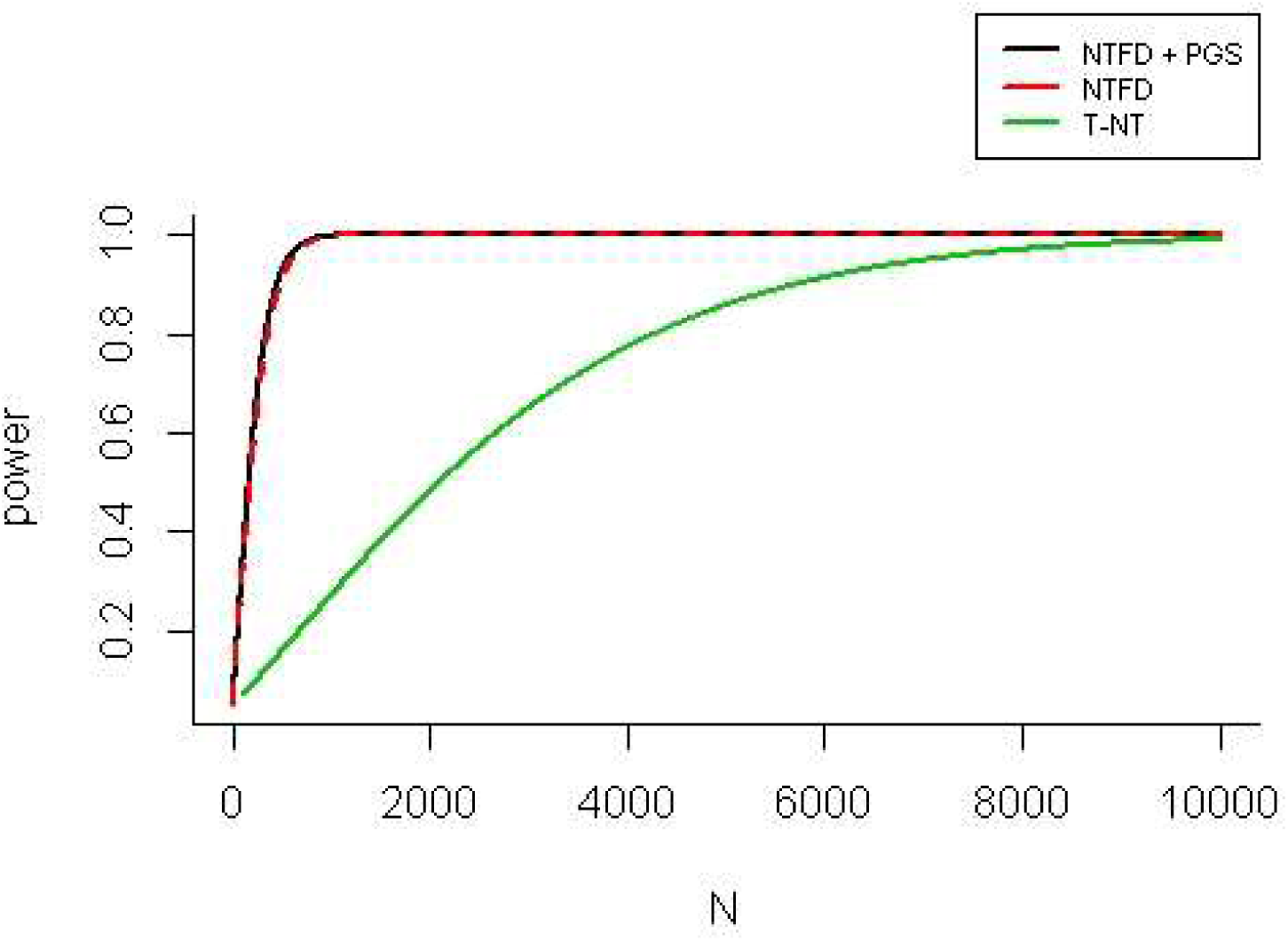
Power plot of the power to detect genotype-environment correlation due to cultural transmission, in three models. This represent scenario 7, where genotype-environment correlation due to cultural transmission explains 15% of the phenotypic variance (2*σ*_*AL,F*_ = .14 and 2*σ*_*PRS,F*_ = .02). in the NTFD designs, N stands for number of families, while in the T-NT design, N is the effective number of unrelated individuals.

As the power of the T-NT design is expected to increase as the additive genetic variance explained by the PRS increases, we tested a model with *a*^2^ = .8, *s*^2^ = .2, *e*^2^ = .5, and *m* = .15, in which the PRS explained either 50% or 100% of the additive genetic variance. We recognize that such high *R*^2^ values are unrealistic; hence this scenario merely serves as a check. As can be seen from Table 3, when the *R*^2^ of the PRS increases, the power to detect cultural transmission also increases, in both the NTFD+PRS and the T-NT design.

**Table 3.**
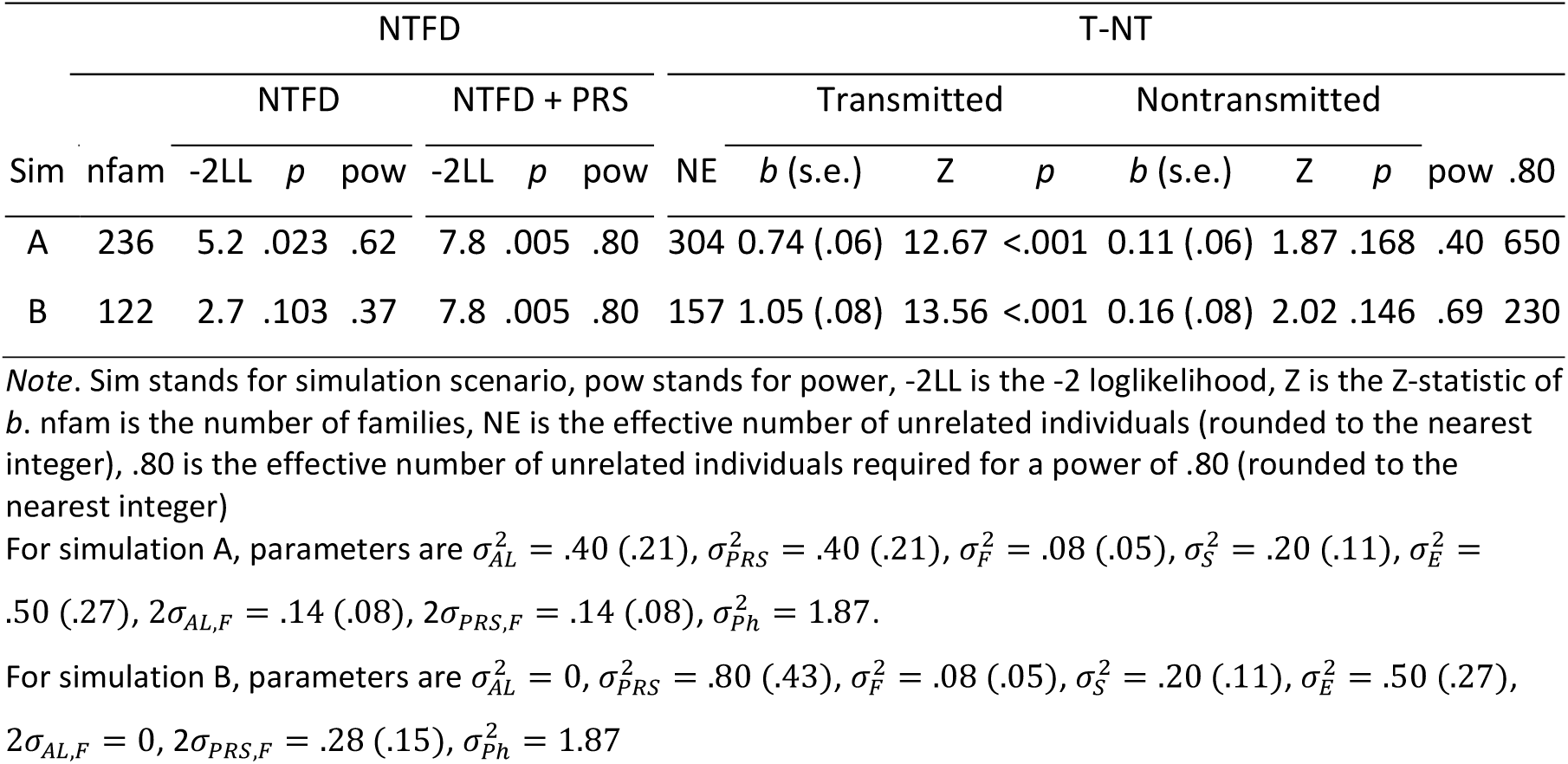
Required sample sizes and power given unstandardized input parameters a^2^ = .8, s^2^ = .2, e^2^ = .5, m = .15 and R^2^ = .5 (sim. A) or R^2^ = 1 (sim B).

## Discussion

The present aim was to compare the power of to detect cultural transmission in the NTF design and the T-NT design. In addition, we explored the possible benefits of incorporating PRS in the NTF design. The classical NTF design is relatively well-powered to detect cultural transmission. However, the inclusion of PRS in the design did not result in an appreciable improvement in power, unless the PRS explain a large portion of the additive genetic variance. Compared to the T-NT design, the NTF design has greater power, given comparable sample sizes and parameter settings. The T-NT design requires much larger samples to detect cultural transmission effects. The difference in power is due to the fact that the transmitted-nontransmitted design, by definition, only captures part of the transmission effects, as the regression of the nontransmitted PRS is an imperfect representation of the total cultural transmission effect. Cultural transmission effects are also conveyed in the transmitted PRS. Currently most PRS only explain a (relatively) small proportion of the total additive genetic variance (e.g., Baselmans et al., 2020). Therefore, the extent to which genotype-environment covariance can be captured by PRSs is proportional to the amount of genetic variance captured by the PRSs. For example, if the total passive genotype-environment covariance accounts for 27% of the phenotypic variance, a PRS explaining 10% of the additive genetic variance will capture only 3% of the phenotypic variance due to the passive genotype-environment covariance (Table 2, simulation 12).

The transmitted-nontransmitted design is an ingenious addition to the designs suitable to detect passive genotype-environment covariance. At present, work is under way to specifying the transmitted-nontransmitted design as a structural equation model (e.g. Bates et al., 2018; Balbona et al, in press; Kim et al., 2020), which will increase its flexibility and scope. For instance, Balbona et al. (2020) and Kim et al (2020) proposed extensions of the transmitted-nontransmitted design in which family environment was modeled as a latent variable. In addition, Kim et al. (2020) proposed that existing bias in the model might be due to unmodeled latent additive genetic variance

In conclusion, the incorporation of PRS in the nuclear twin family design does not provide any benefits to the power to detect cultural transmission, when the polygenic risk scores explain a relatively small part of the total additive genetic variance. The classical nuclear twin family design is itself relatively well-powered to detect cultural transmission, and a decent sample of nuclear twin families is currently more informative with respect to cultural transmission, than is measured genotypic information.

## Supporting information

Rcode

## Acknowledgements

DIB acknowledges Royal Netherlands Academy of Science Professor Award (PAH/6635)

