## Supplementary material for "The power to detect cultural transmission in the Nuclear Twin Family design with and without polygenic risk scores and in the Transmitted-Non-transmitted (alleles) design": Rcode

Supplementary materials

R script

```
#R-scripts for analyses conducted in Bruins, Dolan, & Boomsma.
#settings in correspondence with scenario 4
#other scenarios can be replicated by changing parameter settings and N
#
library(OpenMx)
library(MASS)
#
# a function to calculate power
getchipow=function(alpha,df,Tval) {
  ca=qchisq(alpha,df,ncp=0,lower.tail=F)
  # critical value given alpha
  power=pchisq(ca,df,ncp=Tval,lower.tail=F)
  power
}
#
#
#
#--------------------------------------------------------------------------
#---------------------------------------NTFDs------------------------------
#--------------------------------------------------------------------------
#
# parameter settings for simulation
#
nmz=400  # sample size MZs
ndz=400  # sample size DZz
nfam=nmz+ndz # number of families
#
# ng = n genetic variants, each one has the same effect
# total effect (additive genetic) is equal to a^2, where a is defined below.
#
ng=100  # number of diallelic loci
ngp=10  # number of loci comprising polygenic risk score prs (0 <= npg <= ng).
p_prs=ngp/ng  # percentage explained by prs
p_A=1-p_prs # not explained = A without prs effect
#
# ngp / ng = % explained add gen variance
#
pal=.5     # maf all GVs same maf
qal=1-pal  # major allele freq
#
# symbols from keller 2009 paper
#
s=sqrt(.2)  # shared env twins s^2 = shared (by twins) env variance
a=sqrt(.8)  # add gen a^2 add genetic variance # total A variance
a1=sqrt(a^2*p_A)  # a1 of A (without prs)
a2=sqrt(a^2*p_prs)  # a2 of prs
#
f=sqrt(1)  # constraint scaling - keller. scaling latent variable F (shared env due to m ph->F)
e=sqrt(.5)  # unshared env  e^2 unshared env variance
d=0 # no dominance in this model
m=.10 # parent Ph -> offspring F - cult trans
mu=0 # no phenotypic assort. mating. this version assumes mu=0
q=varA=1 # no ass mating mu=0 variance of A scales to 1
q1=1  # scaling of A without prs
q2=1  # scaling of prs
# derive the value of the A,F covariance
#
w=covAF=(q*a*m) / (1-f*m)  # get w covariance AF: w = m*(q*a+w*f) solve for w
s2=(a^2*q+2*a*w*f+e^2+s^2) / (1-f^2*2*m^2) #get phenotypic variance
x=varF=2*m*s2*m  #variance of the latent variable F
tmpm=matrix(c(a^2,w,w,x),2,2)
xw1=sqrt(a1^2 / a^2)
xw2=sqrt(a2^2 / a^2)
#
w1=covAF1 = (q1*a1*m) / (1-f*m)  # q=1, f=1 covariance A (without prs) and F
w2=covAF2 = (q2*a2*m) / (1-f*m)  # q=1, f=1, covariance A (without prs) and F
#checks (should be 0)
#w1-xw1*w
#w2-xw2*w
w1=xw1*w
w2=xw2*w
#
#Phenotypic variance
s2=(a^2*q+2*a*w*f+e^2+s^2) / (1-f^2*2*m^2) # get phenotypic variance
#varPh = s2
varPh=a^2*q + f^2*x + 2*a*w*f + e^2 + s^2   # eq 1 keller
varPh1=a1^2*q1 + a2^2*q2 + f^2*x + 2*a2*w2*f +2*a1*w1*f + e^2 + s^2   # eq 1 keller
#
# ------------------------end parameter settings---------------------------
#
# exact data simulation
#
# --------preparation for openmx - defining matrices LISREL type model spec
#
ny=8    # t1 t2 M F p1 p2 pM pF
ne=28   # t1 t2 M F p1 p2 pM pF   A D F S E    A D F S E    A D F S E    A D F S E
#
# Ldz: laLdz frLdz vaLdz
# Lmz: laLmz frLmz vaLmz
# Y: laY frY vaY
# B: laB frB vaB
#
cnames=c('t1','t2','M','F','p1','p2','pM','pF','A','D','F','S','E','A','D','F','S','E','A','D','F','S','E','A','D','F','S','E')
rnames=c('t1','t2','M','F','p1','p2','pM','pF','A','D','F','S','E','A','D','F','S','E','A','D','F','S','E','A','D','F','S','E')
#
#Lambda matrix
frL=matrix(FALSE,ny,ne)
vaL=matrix(0,ny,ne); vaL[1:8,1:8]=diag(8)
laL=matrix(NA,ny,ne)
colnames(frL)=cnames
colnames(vaL)=cnames
rownames(vaL)=cnames[1:8]
#
#Beta matrix
vaB=matrix(0,ne,ne)
laB=matrix(NA,ne,ne)
frB=matrix(FALSE,ne,ne)
colnames(frB) = rownames(frB) = cnames
colnames(vaB) = rownames(vaB) = cnames
#
vaB[1:4,5:8]=diag(rep(a2,4))  # prs are standardized so
vaB[1,9:13]=c(a1,d,1,s,e)     # A (without prs), D, F, S, E
vaB[2,14:18]=c(a1,d,1,s,e)
vaB[3,19:23]=c(a1,d,1,s,e)
vaB[4,24:28]=c(a1,d,1,s,e)
vaB[11,3]=vaB[16,3]=m      # Mo -> F (twin1,2)
vaB[11,4]=vaB[16,4]=m      # Fa -> F (twin1,2)
#
vaB[9,19]=vaB[14,19]=vaB[9,24]=vaB[14,24]=.5
vaB[5,7]=vaB[5,8]=vaB[6,7]=vaB[6,8]=.5
#
plab=c('a1','d','f','s','e')
laB[1,5]=laB[2,6]=laB[3,7]=laB[4,8]='a2'
laB[1,9:13]=plab
laB[2,14:18]=plab
laB[3,19:23]=plab
laB[4,24:28]=plab
laB[11,3]=laB[16,3]='m'      # Mo -> F (twin1,2)
laB[11,4]=laB[16,4]='m'      # Fa -> F (twin1,2)
#
frB[1,5]=frB[2,6]=frB[3,7]=frB[4,8]=TRUE  # a2 polygen
frB[1,9:13]=c(TRUE,FALSE,FALSE,TRUE,TRUE)     #  c('a1','d','f','s','e')  d=0 f=1 (fixed)
frB[2,14:18]=c(TRUE,FALSE,FALSE,TRUE,TRUE)    #  c('a1','d','f','s','e')
frB[3,19:23]=c(TRUE,FALSE,FALSE,TRUE,TRUE)    #  c('a1','d','f','s','e')
frB[4,24:28]=c(TRUE,FALSE,FALSE,TRUE,TRUE)    #  c('a1','d','f','s','e')
frB[11,3]=frB[16,3]=TRUE     # Mo -> F (twin1,2)
frB[11,4]=frB[16,4]=TRUE     # Fa -> F (twin1,2)
#
#Psi matrix
vaPs=matrix(0,ne,ne)
laPs=matrix(NA,ne,ne)
frPs=matrix(FALSE,ne,ne)
rownames(vaPs)=cnames
colnames(vaPs)=cnames
frPs[19,21]=frPs[21,19]=frPs[7,21]=frPs[21,7]=TRUE
frPs[24,26]=frPs[26,24]=frPs[8,26]=frPs[26,8]=TRUE
frPs[21,21]=frPs[26,26]=TRUE
#
laPs[19,21]=laPs[21,19]='w1'
laPs[7,21]=laPs[21,7]='w2'
laPs[24,26]=laPs[26,24]='w1'
laPs[8,26]=laPs[26,8]='w2'
laPs[21,21]=laPs[26,26]='x'
#
diag(vaPs) = 1
vaPs[5,5]=vaPs[6,6]=vaPs[9,9]=vaPs[14,14]=.5 # residual A variance
diag(vaPs[1:4,1:4])=0  # phenotypes zero residual variance : all variance explained by model
#
vaPs[11,11]=vaPs[16,16]=0  #no residual F in twins
vaPs[19,21]=vaPs[21,19]=w1 #cov(A,F)
vaPs[7,21]=vaPs[21,7]=w2 #cov(PRS,F)
vaPs[24,26]=vaPs[26,24]=w1 #cov(A,F)
vaPs[8,26]=vaPs[26,8]=w2 #cov(PRS,F)
vaPs[21,21]=vaPs[26,26]=x  # variance of latent variable F
#
vaPs[12,17]=vaPs[17,12]=1  #S
vaPs[10,15]=vaPs[15,10]=.25  # D
#
# DZ families model Phenotype + PRS
#
Id=diag(ne) # ne=28
IBi=solve(Id-vaB)
tmp=IBi%*%vaPs%*%t(IBi)
Sdz1=vaL%*%IBi%*%vaPs%*%t(IBi)%*%t(vaL)
#
# Get MZ families model Pheno + prs ... via Ps: cor(D,D)=1 residual A cov =.5
vaPsm=vaPs
vaPsm[5,6]=vaPsm[6,5]=vaPsm[9,14]=vaPsm[14,9]=.5  # A and prs in MZ
vaPsm[10,15]=vaPsm[15,10]=1  # D in MZ
#
Smz1=vaL%*%IBi%*%vaPsm%*%t(IBi)%*%t(vaL)
#
#Smz1 and Sdz1 are the expected exact covariance matrices.
#
# ----------------------------------------------exact data sim---------------------------------
#
ephdatmz=mvrnorm(nmz, mu=rep(0,8), Sigma=Smz1, emp=TRUE)
ephdatdz=mvrnorm(nmz, mu=rep(0,8), Sigma=Sdz1, emp=TRUE)
vnames2=c('T1','T2','Mo','Fa','pg1','pg2','pgm','prs')
colnames(ephdatmz)=vnames2
colnames(ephdatdz)=vnames2
ephdatmz=as.data.frame(ephdatmz)
ephdatdz=as.data.frame(ephdatdz)
#
# ----------------------------end exact data sim---------------------------
#
#  start model fitting 1) full model exact
#
ny=8    # t1 t2 M F p1 p2 pM pF
ne=28   # t1 t2 M F p1 p2 pM pF   A D F S E    A D F S E    A D F S E    A D F S E
#
# starting values... change to check
dost=F
if (dost) {
  a2=.2
  a1=.6
  s=.05
  e=.6
  m=.13
  w1=.1
  w2=.05
  x=.2
  # change starting values in matrices
  vaB[1:4,5:8]=diag(rep(a2,4))  # prs are standardized so
  vaB[1,9:13]=c(a1,d,1,s,e)     # A (without prs), D, F, S, E
  vaB[2,14:18]=c(a1,d,1,s,e)
  vaB[3,19:23]=c(a1,d,1,s,e)
  vaB[4,24:28]=c(a1,d,1,s,e)
  vaB[11,3]=vaB[16,3]=m      # Mo -> F (twin1,2)
  vaB[11,4]=vaB[16,4]=m      # Fa -> F (twin1,2)
  vaPs[19,21]=vaPs[21,19]=w1
  vaPs[7,21]=vaPs[21,7]=w2
  vaPs[24,26]=vaPs[26,24]=w1
  vaPs[8,26]=vaPs[26,8]=w2
  vaPs[21,21]=vaPs[26,26]=x  # variance of x
  vaPsm=vaPs  # mz matrix
  vaPsm[5,6]=vaPsm[6,5]=vaPsm[9,14]=vaPsm[14,9]=.5  # A and prs in MZ
  vaPsm[10,15]=vaPsm[15,10]=1  # D in MZ
}
#end starting values #
#openmx 1
#first fit the model to the phenotypic data only.
ephdatmz0=ephdatmz[,c(1,2,3,4)]
ephdatdz0=ephdatdz[,c(1,2,3,4)]
vnames=colnames(ephdatmz0)    # t1 t2 m f
#
#------openMx based on Keller (2009) script ---- standard model without prs
#
ntf <- mxModel(model="NTF",
               # Matrices
               mxMatrix(type="Full", nrow=1, ncol=1, free=T, values=.2,  label="FamilialPath", name="m"),
               # fix m=0 if you want Vf=0
               mxMatrix(type="Full", nrow=1, ncol=1, free=TRUE, values=.2,  label="Sib",          name="s"),
               # fix s=0 if you want Vs=0  # cvd; Vs is zero
               mxMatrix(type="Full", nrow=1, ncol=1, free=TRUE, values=.4,  label="Env",          name="e"),
               mxMatrix(type="Full", nrow=1, ncol=1, free=TRUE, values=.4,  label="AddGen",       name="a"),
               # cvd:D out
               mxMatrix(type="Full", nrow=1, ncol=1, free=FALSE, values=.0,  label="Dominance",    name="d"),
               # D not used or used
               mxMatrix(type="Full", nrow=1, ncol=1, free=FALSE, values=.0,  label="AMCopath",     name="mu"),
               mxMatrix(type="Full", nrow=1, ncol=1, free=TRUE, values=1, label="VarPhen",      name="Vp1"),
               mxMatrix(type="Full", nrow=1, ncol=1, free=T, values=.1,  label="VarF",         name="x1"),
               # keep this parameter free, even if Vf is fixed to 0
               mxMatrix(type="Full", nrow=1, ncol=1, free=T, values=.2, label="CovPhenGen",   name="delta1"),
               mxMatrix(type="Full", nrow=1, ncol=1, free=FALSE, values=1,   label="VarAddGen",    name="q1"),
               mxMatrix(type="Full", nrow=1, ncol=1, free=T, values=.1, label="CovFA",        name="w1"),
               #
               # f = 1 fixed identifying constraint (keller)
               #
               mxMatrix(type="Full", nrow=1, ncol=1, free=FALSE, values=1, label="fpath",name="f"),
               # fix it to 1 or to true value (in simulation)
               # mxAlgebra section - nonlinear constraints
               mxAlgebra(e %*% t(e), name="E"),
               mxAlgebra(d %*% t(d), name="D"),
               mxAlgebra(s %*% t(s), name="S"),
               #
               mxAlgebra((a%*%a%*%q1) + (f%*%f%*%x1) + 2%x%a%*%w1%*%f + E + D + S, name="Vp2"),
               # cvd:
               mxAlgebra( 2%x%(m%*%Vp1%*%m) + 2%x%(m%*%Vp1%*%mu%*%Vp1%*%m) , name="x2"),
               mxAlgebra(q1%*%a + w1%*%f, name="delta2"),
               mxAlgebra(1 + delta1 %*% mu %*% t(delta1), name="q2"),
               mxAlgebra(delta1 %*% m + delta1 %*% mu %*% Vp1 %*% m, name="w2"),
               #
               #constraints - equating nonlinear constraints and parameters
               #
               mxConstraint(Vp1==Vp2,   name='VpCon'),
               mxConstraint(x1==x2,    name='xCon'),
               mxConstraint(delta1==delta2,name='deltaCon'),
               mxConstraint(q1==q2,    name='qCon'),
               mxConstraint(w1==w2,    name='wCon'),
               #mxAlgebra section - relative covariances
               mxAlgebra((a%*%a%*%q1) + (f%*%f%*%x1) + 2%x%(a%*%w1%*%f) + D + S, name="CvMz"),
               mxAlgebra((a%*%a%*%(q1-.5))  + .25%x%D + (f%*%f%*%x1) + 2%x%(a%*%w1%*%f) + S, name="CvDz"),
               #
               mxAlgebra( .5%x%a%*%(q1%*%a + w1%*%f) + .5%x%a%*%(q1%*%a + w1%*%f)%*%mu%*%Vp1 +
                            m%*%Vp1 + m%*%(Vp1%*%Vp1%*%mu), name="ParChild"),
               mxAlgebra(Vp1%*%Vp1%*%mu,name="CvSps")
)
#
# #  MZ group ----------------------------------------------------
mzModel <- mxModel(name = "MZNTF",
                   mxMatrix(type="Full", nrow=1, ncol=4, free=TRUE, values= .0, label="mean", name="expMeanMz"),
                   # Algebra for expected variance/covariance matrix in MZF
                   mxAlgebra(expression=rbind(
                     cbind(NTF.Vp1,      NTF.CvMz,     NTF.ParChild, NTF.ParChild),
                     cbind(NTF.CvMz,     NTF.Vp1,      NTF.ParChild, NTF.ParChild),
                     cbind(NTF.ParChild, NTF.ParChild, NTF.Vp1,      NTF.CvSps),
                     cbind(NTF.ParChild, NTF.ParChild, NTF.CvSps,    NTF.Vp1) ),
                     dimnames=list(vnames,vnames),name="expCovMz"),
                   mxData(observed=ephdatmz0, type="raw"),
                   mxExpectationNormal(covariance="expCovMz", means = "expMeanMz", vnames),
                   mxFitFunctionML() )
#
# DZ group -------------------------------------------------------------------
dzModel <- mxModel(name = "DZNTF",
                   mxMatrix(type="Full", nrow=1, ncol=4, free=TRUE, values= .0, label="mean", name="expMeanDz"),
                   # Algebra for expected variance/covariance matrix in MZF
                   mxAlgebra(expression=rbind(
                     cbind(NTF.Vp1,      NTF.CvDz,     NTF.ParChild, NTF.ParChild),
                     cbind(NTF.CvDz,     NTF.Vp1,      NTF.ParChild, NTF.ParChild),
                     cbind(NTF.ParChild, NTF.ParChild, NTF.Vp1,      NTF.CvSps),
                     cbind(NTF.ParChild, NTF.ParChild, NTF.CvSps,    NTF.Vp1) ),
                     dimnames=list(vnames,vnames),name="expCovDz"),
                   mxData(observed=ephdatdz0, type="raw"),
                   mxExpectationNormal(covariance="expCovDz", means = "expMeanDz", vnames),
                   mxFitFunctionML() )
#
# assemble the model
NTFModel0 <- mxModel(model="NucTwFam", mzModel, dzModel, ntf,
                     mxFitFunctionMultigroup( c("MZNTF","DZNTF") )  )
#
#Run MX ---- run the model using NPSOL
#
mxOption(NULL, "Default optimizer"  , "NPSOL")
NTFModelFit0 <- mxRun(NTFModel0)
#
# --------------------------- second
# --------------------------- NTF + PRS design
#
ntf2 <- mxModel(model="NTF2",
                 # LISREL type
                 mxMatrix(type="Full", nrow=ny, ncol=ne, free=frL, values=vaL, label=laL, name="Ly"),
                 mxMatrix(type="Full", nrow=ne, ncol=ne, free=frB, values=vaB, label=laB,name="Be"),
                 mxMatrix(type="Symm", nrow=ne, ncol=ne, free=frPs, values=vaPs,  label=laPs,name="Psd"),
                 mxMatrix(type="Symm", nrow=ne, ncol=ne, free=frPs, values=vaPsm,  label=laPs,name="Psm"),
                 mxMatrix(type="Diag", nrow=ne, ncol=ne, free=FALSE, values=1,  label=NA,name="Id"),
                 mxMatrix(type="Diag", nrow=1, ncol=1, free=FALSE, values=1,  label=NA,name="Id1"),
                 # parameters
                 mxMatrix(type="Full", nrow=1, ncol=1, free=T, values=w1, label='w1', name="W1"),
                 mxMatrix(type="Full", nrow=1, ncol=1, free=T, values=w2, label='w2', name="W2"),
                 mxMatrix(type="Full", nrow=1, ncol=1, free=T, values=m, label='m', name="M"),
                 mxMatrix(type="Full", nrow=1, ncol=1, free=T, values=1, label='sph1', name='SPH1'),
                 mxMatrix(type="Full", nrow=1, ncol=1, free=T, values=a1, label='a1', name='A1'),
                 mxMatrix(type="Full", nrow=1, ncol=1, free=T, values=a2, label='a2', name='A2'),
                 mxMatrix(type="Full", nrow=1, ncol=1, free=T, values=x, label='x', name='X'),
                 mxMatrix(type="Full", nrow=1, ncol=1, free=T, values=e, label='e', name='E'),
                 mxMatrix(type="Full", nrow=1, ncol=1, free=F, values=.0, label='d', name='D'),
                 mxMatrix(type="Full", nrow=1, ncol=1, free=T, values=s, label='s', name='S'),
                 #
                 mxAlgebra((A1%*%A1+A2%*%A2),   name='A12'),
                 mxAlgebra(sqrt((A1%*%A1)/A12),   name='r1'),
                 mxAlgebra(sqrt((A2%*%A2)/A12),   name='r2'),
                 mxAlgebra(((sqrt(A12)%*%M)%*%solve(Id1-M)),   name='W'),
                 mxConstraint(W1==r1%*%(W),   name='w1Con'),
                 mxConstraint(W2==r2%*%(W),   name='w2Con'),
                 mxConstraint(X==(2%x%(M%*%SPH1%*%M)),   name='xCon'),
                 mxConstraint(SPH1==A1%*%A1+A2%*%A2+X+2%x%(A1%*%W1)+2%x%(A2%*%W2)+E%*%E+S%*%S+D%*%D, name='s2Con'),
                 #
                 mxAlgebra(expression = solve(Id-Be), name='IBi'),
                 mxAlgebra(expression = Ly%*%IBi, name='LyIBi'),
                 mxAlgebra(expression = LyIBi%*%Psd%*%t(LyIBi), name='Sdz'),
                 mxAlgebra(expression = (LyIBi%*%Psm%*%t(LyIBi))[-6,-6], name='Smz')
)
#
dzModel2 <- mxModel(name = "DZNTF2",
                     mxMatrix(type="Full", nrow=1, ncol=8, free=TRUE, values= .0,
                              label=c("phm","phm","phm","phm","pm","pm","pm","pm"), name="expMeanDz"),
                     # Algebra for expected variance/covariance matrix in MZF
                     mxData(observed=ephdatdz, type="raw"),
                     mxExpectationNormal(covariance="NTF2.Sdz", means = "expMeanDz", vnames2),
                     mxFitFunctionML() )
#
mzModel2 <- mxModel(name = "MZNTF2",
                     mxMatrix(type="Full", nrow=1, ncol=7, free=TRUE, values= .0,
                              label=c("phm","phm","phm","phm","pm","pm","pm"), name="expMeanMz"),
                     # Algebra for expected variance/covariance matrix in MZF
                     mxData(observed=ephdatmz[,-6], type="raw"),
                     mxExpectationNormal(covariance="NTF2.Smz", means = "expMeanMz", vnames2[-6]),
                     mxFitFunctionML() )
#
# assemble the model
NTFModel2<- mxModel(model="NucTwFam2", mzModel2, dzModel2, ntf2,
                     mxFitFunctionMultigroup( c("MZNTF2","DZNTF2") )  )
#
NTFModelFit2 <- mxRun(NTFModel2)
#
###################################
# ------------------------------- end full model exact data
summary(NTFModelFit0) #keller
summary(NTFModelFit2) #NTD+PRS
round(coef(NTFModelFit0),2)
round(coef(NTFModelFit2),2)
#
#----------------------------------Power Analysis
#----------------------------------Model comparisons to obtain NPC
NTFModel0_m = omxSetParameters(NTFModel0, labels='FamilialPath', value=0, free=F)
NTFModelFit0_m <- mxRun(NTFModel0_m)
t0_m=mxCompare(NTFModelFit0, NTFModelFit0_m)
NCP0=t0_m[2,7]  # this is the ratio, but given exact data sim this is the NCP
NTFModel2_m = omxSetParameters(NTFModel2, labels='m', value=0, free=F)
NTFModelFit2_m <- mxRun(NTFModel2_m)
t2_m=mxCompare(NTFModelFit2, NTFModelFit2_m)
NCP2=t2_m[2,7]  # this is the ratio, but given exact data sim this is the NCP
#
#
alpha=.05
df=1
print(getchipow(alpha, 1, NCP0)) #NTFD power
print(getchipow(alpha, 1, NCP2)) #NTFD+PRS power
#
#
#--------------------------------------------------------------------------
#--------------------------------------TNT---------------------------------
#--------------------------------------------------------------------------
#
#
library(MASS)
library(gee)
#
nfam=800 #NTFD+PGD had a power of .80 with nfam=800, now run TNT design with same parameter settings
#
Nrep=5000
N=100
parametergee<-matrix(0,Nrep,9) #store parameters here
alpha=0.05
#
result=matrix(0,Nrep,2)
#
powerTNT<-function(result){
  set.seed(1)
  #same specifications as before in NTFD+PRS
  nmz=nfam/2
  ndz=nfam/2
  ng=100
  ngp=10
  p_prs=ngp/ng
  p_A=1-p_prs
  pal=.5
  qal=1-pal
  s=sqrt(.2)
  a=sqrt(.8)
  a1=sqrt(a^2*p_A)
  a2=sqrt(a^2*p_prs)
  f=sqrt(1)
  e=sqrt(.5)
  d=0
  m=.1
  mu=0
  q=varA=1
  q1=1
  q2=1
  w=covAF=(q*a*m) / (1-f*m)
  s2=(a^2*q+2*a*w*f+e^2+s^2) / (1-f^2*2*m^2)
  x=varF=2*m*s2*m
  tmpm=matrix(c(a^2,w,w,x),2,2)
  xw1=sqrt(a1^2 / a^2)
  xw2=sqrt(a2^2 / a^2)
  w1=xw1*w
  w2=xw2*w
  #
  varPh=a^2*q + f^2*x + 2*a*w*f + e^2 + s^2
  #
  #---------------------------------------end parameter settings
  #---------------------------------------data simulation
  Nrep=5000
  N=100
  alpha=0.05
  result=matrix(0,Nrep,2)
  for(reps in 1:Nrep){
    # simulate parental alleles assuming random mating and linkage equilibrium
    am=array(0,c(nfam,ng,2)) # mother alleles
    af=array(0,c(nfam,ng,2)) # father alleles
    gm=matrix(0,nfam,ng) # mother genotype
    gf=matrix(0,nfam,ng) # father genotype
    Am=matrix(0,nfam,1)  # mother prs
    Af=matrix(0,nfam,1)  # father prs
    prsm=matrix(0,nfam,1) # mother polygenic scores
    prsf=matrix(0,nfam,1) # father polygenic scores
    #
    at1=array(0,c(nfam,ng,2))  # twin 1 alleles transmitted from m and f
    at2=array(0,c(nfam,ng,2))  # twin 2 alleles transmitted from m and f
    ant1=array(0,c(nfam,ng,2))  # t1 alleles not transmitted - ibd = 0 with t1
    ant2=array(0,c(nfam,ng,2))  # t2 alleles not transmitted - ibd = 0 with t2
    g1=matrix(0,nfam,ng)  # genotypes twin 1
    g2=matrix(0,nfam,ng)  # genotypes twin 2
    A1=matrix(0,nfam,2) # prs tw 1 based on 1) transmitted, 2) based on non-transmitted
    A2=matrix(0,nfam,2) # prs tw 2 based on 1) transmitted, 2) based on non-transmitted
    prs1=matrix(0,nfam,2) # twin 1 polygenic scores 1=transmitted 2=not transmitted
    prs2=matrix(0,nfam,2) # twin 2 polygenic scores 1=transmitted 2=not transmitted
    #
    # parental alleles
    #
    for (i in 1:ng) {
      am[,i,1] = sample(c(0,1),nfam,replace=T,prob=c(pal,qal))
      am[,i,2] = sample(c(0,1),nfam,replace=T,prob=c(pal,qal))
      af[,i,1] = sample(c(0,1),nfam,replace=T,prob=c(pal,qal))
      af[,i,2] = sample(c(0,1),nfam,replace=T,prob=c(pal,qal))
    }
    #
    for (i in 1:ng) {
      # twin 1 inherits alleles from father and mother
      mt1=sample(c(1,2),nfam,replace=T,prob=c(.5,.5)) # sample maternal alleles for transmission
      ft1=sample(c(1,2),nfam,replace=T,prob=c(.5,.5)) # sample paternal alleles for transmission
      mnt1=3-1*mt1  # 1->2 2-> 1  # nontransmitted maternal alleles
      fnt1=3-1*ft1 # 1->2 2-> 1  # nontransmitted paternal alleles
      # twin 2
      mt2=sample(c(1,2),nfam,replace=T,prob=c(.5,.5))
      ft2=sample(c(1,2),nfam,replace=T,prob=c(.5,.5))
      mnt2=3-1*mt2
      fnt2=3-1*ft2
      # offsprin alleles transmitted and not transmitted
      for (k in 1:nfam) {
        at1[k,i,1] = am[k,i,mt1[k]]   # transm
        at1[k,i,2] = af[k,i,ft1[k]]   # transm
        ant1[k,i,1]= am[k,i,mnt1[k]]  # not transm
        ant1[k,i,2]= af[k,i,fnt1[k]]  # not transm
        at2[k,i,1] = am[k,i,mt2[k]]   # transm
        at2[k,i,2] = af[k,i,ft2[k]]   # transm
        ant2[k,i,1]= am[k,i,mnt2[k]]  # not transm
        ant2[k,i,2]= af[k,i,fnt2[k]]  # not transm
      } # nfam
    } # ng
    # A scores polygenic scores representing A = total add gen
    for (i in 1:ng) {
      Am[,1]=Am[,1]+(am[,i,1]+am[,i,2])
      Af[,1]=Af[,1]+(af[,i,1]+af[,i,2])
      A1[,1]=A1[,1]+(at1[,i,1]+at1[,i,2])
      A1[,2]=A1[,2]+(ant1[,i,1]+ant1[,i,2])
      A2[,1]=A2[,1]+(at2[,i,1]+at2[,i,2])
      A2[,2]=A2[,2]+(ant2[,i,1]+ant2[,i,2])
    }
    #
    # PRS based on first ngp genetic variants
    #
    for (i in 1:ngp) {
      prsm[,1]=prsm[,1]+(am[,i,1]+am[,i,2])
      prsf[,1]=prsf[,1]+(af[,i,1]+af[,i,2])
      prs1[,1]=prs1[,1]+(at1[,i,1]+at1[,i,2])
      prs1[,2]=prs1[,2]+(ant1[,i,1]+ant1[,i,2])
      prs2[,1]=prs2[,1]+(at2[,i,1]+at2[,i,2])
      prs2[,2]=prs2[,2]+(ant2[,i,1]+ant2[,i,2])
    }
    #
    # scale A to variance = 1  parameter a is the effect
    #
    Am[,1]=scale(Am[,1])   # polygenic score mother standardized
    Af[,1]=scale(Af[,1])   # polygenic score father standardized
    A1[,1]=scale(A1[,1])   # polygenic score tw1 standardized (transmitted alleles)
    A1[,2]=scale(A1[,2])   # polygenic score tw1 standardized (not transmitted alleles)
    A2[,1]=scale(A2[,1])   # polygenic score tw2 standardized (transmitted alleles)
    A2[,2]=scale(A2[,2])   # polygenic score tw2 standardized (not transmitted alleles)
    #
    # scale polygenic scores
    #
    prsm[,1]=scale(prsm[,1])   # polygenic score mother standardized
    prsf[,1]=scale(prsf[,1])   # polygenic score father standardized
    prs1[,1]=scale(prs1[,1])   # polygenic score tw1 standardized (transmitted alleles)
    prs1[,2]=scale(prs1[,2])   # polygenic score tw1 standardized (not transmitted alleles)
    prs2[,1]=scale(prs2[,1])   # polygenic score tw2 standardized (transmitted alleles)
    prs2[,2]=scale(prs2[,2])   # polygenic score tw2 standardized (not transmitted alleles)
    #
    Sm=rnorm(nfam,0,1)  # mother C shared by twins but unshared by spouses
    Sf=rnorm(nfam,0,1)  # father C  shared by twins but unshared by spouses
    Em=rnorm(nfam,0,1)  # mother E unshared
    Ef=rnorm(nfam,0,1)  # father E unshared
    #
    # create F so that the covariance with A is correct (covAF) and the variance of F equals varF
    # based on cholesky decomp.
    #
    Fm=Am[1:nfam,1]*covAF + rnorm(nfam,0,1)*sqrt(varF - covAF^2)
    Ff=Af[1:nfam,1]*covAF + rnorm(nfam,0,1)*sqrt(varF - covAF^2)
    #
    #  S & E in twins
    #
    S1=rnorm(nfam,0,1)    # shared E by twins
    S2=S1  # correlated 1  # shared by twins
    E1=rnorm(nfam,0,1)  #   unshared E
    E2=rnorm(nfam,0,1)  #   unshared E
    #
    phm=s*Sm+a*Am[,1]+f*Fm+e*Em   # mother pheno
    phf=s*Sf+a*Af[,1]+f*Ff+e*Ef   # father pheno
    #
    F1=m*phm + m*phf    # parental pheno to F in twins
    F2=m*phm + m*phf    # cult transmission
    #
    # dz
    #
    ph1=s*S1 + a*A1[,1] + f*F1 + e*E1  # offspring 1 pheno
    ph2=s*S2 + a*A2[,1] + f*F2 + e*E2  # offspring 2 pheno
    #
    # 1:ndz  first half
    #
    phdatdz=matrix(0,ndz,4)
    phdatdz[,1]=phm[1:ndz]
    phdatdz[,2]=phf[1:ndz]
    phdatdz[,3]=ph1[1:ndz]
    phdatdz[,4]=ph2[1:ndz]
    cov(phdatdz)
    #
    # mz
    #
    ph1=s*S1 + A1[,1]*a + f*F1 + e*E1
    ph2=s*S2 + A1[,1]*a + f*F2 + e*E2  # mz genetically identical A1 twin1 = A1 twin2
    # ndz+1 : nfam second half
    phdatmz=matrix(0,nmz,4)
    phdatmz[,1]=phm[(ndz+1):nfam]
    phdatmz[,2]=phf[(ndz+1):nfam]
    phdatmz[,3]=ph1[(ndz+1):nfam]
    phdatmz[,4]=ph2[(ndz+1):nfam]
    #
    # the expected covariance matrices based on the parameters
    #
    ESdz=matrix(0,4,4)
    ESdz[1,1]=ESdz[2,2]=ESdz[3,3]=ESdz[4,4]=varPh
    ESdz[1,2]=ESdz[2,1]=mu # zero
    #                                                 zero...................             zero............
    ESdz[3,1]=ESdz[3,2]=ESdz[4,1]=ESdz[4,2]= .5*a*(q*a+w*f) + .5*a*(q*a+w*f)*mu*varPh + m*varPh + m*varPh*mu*varPh
    ESdz[1,3]=ESdz[2,3]=ESdz[1,4]=ESdz[2,4]= .5*a*(q*a+w*f) + .5*a*(q*a+w*f)*mu*varPh + m*varPh + m*varPh*mu*varPh
    ESdz[4,3]=ESdz[3,4]=a^2*(q-.5)+s^2+f^2*x+2*a*w*f
    #
    ESmz=ESdz
    ESmz[4,3]=ESmz[3,4]=a^2*q+s^2+f^2*x+2*a*w*f
    #
    # phdatmz ... add polygenic scores
    #                 mother     father      dz1 t, nt   dz2 t,nt
    pgdatdz=cbind(prsm[1:ndz,],prsf[1:ndz,],prs1[1:ndz,],prs2[1:ndz,])
    #                 mother            father                  mz1 t, nt =   mz2 t, nt (duplicate 1)
    pgdatmz=cbind(prsm[(1+ndz):nfam,],prsf[(1+ndz):nfam,],prs1[(1+ndz):nfam,],prs1[(1+ndz):nfam,])
    #
    phdatmz=as.data.frame(cbind(phdatmz,pgdatmz))
    phdatdz=as.data.frame(cbind(phdatdz,pgdatdz))
    #
    colnames(phdatdz) = colnames(phdatmz) =  vnames1 =
      c('phm','phf','pht1','pht2',
        'prsm','prsf','prst1','prsnt1','prst2','prsnt2')
    #
    # phdatmz and phdatdz are the raw simulated dataset. the order: ph: m fm t1 t2 and prs: m f t1 t2
    # # -------------------------------------------------- N NT simulation
    #                         ph ph p1 p2 p1 p2
    #                         1   2  3  4  5  6
    phdatmzdz=rbind(phdatmz[,c(3,4,7,8,9,10)], phdatdz[,c(3,4,7,8,9,10)])
    info=cbind(c(1:nfam), c(rep(1,nmz),rep(2,ndz)))
    colnames(info)=c('famnr','zyg')
    phdatmzdz=cbind(phdatmzdz,info)
    #                1     2    3    4    5
    # long format famnr, ppnr, ph, prst, prsnt
    indi=matrix(c(
      3,4,
      5,6),2,2,byrow=T)
    Lphdatmzdz=matrix(0,nfam*2,6)
    Lphdatmzdz[,6]=2
    Lphdatmzdz[1:(nmz*2)]=1   # mzs
    ii=0
    for (i in 1:nfam) {
      for (j in 1:2) {
        ii=ii+1
        Lphdatmzdz[ii,1]=i
        Lphdatmzdz[ii,2]=j
        Lphdatmzdz[ii,3]=phdatmzdz[i,j]  # j = 1,2
        k1=indi[j,1]
        k2=indi[j,2]
        Lphdatmzdz[ii,4]=phdatmzdz[i,k1]  # k1 = 3 or 5
        Lphdatmzdz[ii,5]=phdatmzdz[i,k2]  # k2 = 4 or 6
      }}
    colnames(Lphdatmzdz) = c('famnr','pp','ph','pt','pnt','zyg')
    Lphdatmzdz=as.data.frame(Lphdatmzdz)
    #
    tnt1<-gee(ph~pt+pnt, id=famnr, corstr='exchangeable', data=Lphdatmzdz,silent = T)
    sr1=summary(tnt1)
    testst=sr1$coefficients[3,5]^2  # is a z but z^2 is chi2. - robust Z of nt
    pts=2*pnorm(abs(sr1$coefficients[3,5]), lower.tail = FALSE)  #SB: GEE does not give pval, only a Zstat.
    result[reps,1]<<-testst - 1
    result[reps,2]<<-as.numeric((pts<alpha))
    #
    parametergee[reps,1]<<-sr1$coefficients[1,1] #b0
    parametergee[reps,2]<<-sr1$coefficients[2,1] #b1
    parametergee[reps,3]<<-sr1$coefficients[3,1] #b2
    parametergee[reps,4]<<-sr1$coefficients[2,5] #rob. z b1 (pt)
    parametergee[reps,5]<<-sr1$coefficients[2,4] #rob. SE b1
    parametergee[reps,6]<<-sr1$coefficients[3,5] #rob. Z b2 (pnt)
    parametergee[reps,7]<<-sr1$coefficients[3,4] #rob. z b2
    parametergee[reps,8]<<-2*pnorm(abs(sr1$coefficients[2,5]), lower.tail = FALSE) #p b1
    parametergee[reps,9]<<-2*pnorm(abs(sr1$coefficients[3,5]), lower.tail = FALSE) #p b2
    #
  }
  nEff<<-(2*nmz/(1+(ESmz[3,4]/ESmz[3,3])))+(2*ndz/(1+(ESdz[3,4]/ESdz[3,3])))
}
powerTNT(result)
#
#  b0   b1   b2   z(b1) SE (b1) z(b2) se(b2) p(b1) p(b2)
round(apply(parametergee, 2, mean),3) #parameters
apply(result,2,mean)    # mean NCP and power
```
